# Deep learning-based counting and body-area measurement enable sensitive population-level endpoints in the Collembola reproduction test

**DOI:** 10.64898/2026.10.19.760133

**Authors:** Hyun-Gi Min, Lee-Hyeon Jeon, Jeongwon Choi, Han Soo Kim, June Wee, Wonjun Hong, Daejin Choi, Yun-Sik Lee

**Author notes:** **Corresponding author:** Yun-Sik Lee, Daejin Choi. These authors contributed equally to this work as first authors (Hyun-Gi Min, Lee-Hyeon Jeon).

## Abstract

Springtails (Collembola) are standard test organisms for soil ecotoxicological assessment; under the ISO 11267 and OECD 232 guidelines, adult abundance and juvenile production are the primary endpoints. However, these rely on labor-intensive manual counting and capture only abundance, overlooking sublethal effects on growth. We developed an automated system that couples YOLOv8 Object detection with Voronoi–Otsu segmentation-based body-area measurement, so that adults and juveniles of *Allonychiurus kimi* are counted and their growth (body area) measured simultaneously, and applied it to toxicity assessment based on average and cumulative body area and to cohort analysis of juvenile body-area distributions using a Gaussian mixture model (GMM). The model was trained on 142 annotated images and validated on 160 images from toxicity tests with cadmium (Cd), arsenic (As), and copper (Cu), and both detection accuracy and agreement with manual measurements were excellent (coefficient of determination, *R*^2^ ≥ 0.95). Cumulative body area—a composite indicator integrating abundance and individual size, and thus reflecting both reproduction and growth—showed the lowest EC_50_ in all three metal(loid)s (Cd 14.10, As 10.53, Cu 398.80 mg kg^−1^). Cohort analysis showed that the two cohorts present in the controls merged into one at the highest concentrations (As 20, Cu 800 mg kg^−1^), indicating that toxic stress drives not only individual-level suppression of reproduction and growth but also population restructuring. Beyond accurately and efficiently replacing manual counting, the system adds more sensitive composite endpoints and population-level analysis, providing a more comprehensive, scalable framework for soil ecotoxicological risk assessment.

## 1. Introduction

Soil ecotoxicity assessment plays a central role in evaluating the potential risks that chemical pollutants such as metal(loid)s pose to terrestrial ecosystems (van Gestel, 2012). The effect concentrations derived from these assessments underpin soil quality criteria and the ecological risk assessment of contaminated land, so the sensitivity and ecological realism of the endpoints used directly determine how protective those criteria are (Joimel et al., 2022). Early assessments relied on acute toxicity tests focused on survival, but as survival alone proved insufficient to reflect pollutant effects, the range of endpoints was expanded to sublethal indicators such as reproduction and growth (OECD, 1984; ISO, 1999). More recently, biomarker- and omics-based molecular approaches have been proposed as early-warning indicators (Spurgeon et al., 2005; Van Straalen and Feder, 2012), advancing the field toward a more precise, multi-tiered framework. Among the organisms used within this framework, Collembola are one of the most abundant arthropod groups in soils worldwide (Fountain and Hopkin, 2005). They perform key ecosystem functions—including organic matter decomposition, regulation of fungal communities, and soil carbon and nitrogen cycling (Filser, 2002)—and their high sensitivity to diverse pollutants (Joimel et al., 2022) has made them widely used indicator organisms, assessed through endpoints such as survival, reproduction, and avoidance behavior. These characteristics have led to their official adoption in international standard protocols such as ISO 11267 and OECD TG 232.

Both standard protocols designate survival and reproduction (the number of juveniles produced) as core endpoints. Growth, although not an official endpoint, has been recognized as an important complementary indicator of population-level effects, given its trade-off with reproduction (van Gestel, 2012). Studies using growth indicators fall into two categories. The first tracks growth in inoculated adults, detecting toxic responses within a shorter period than the standard reproduction test and at concentrations below those affecting survival (Gruss et al., 2022, 2024). The second analyzes the distribution of growth indicators among the juveniles produced, resolving changes in population structure and intergenerational effects that size-class categorization or reproduction counts alone cannot reveal (Campiche et al., 2007; Marks et al., 2015; Alves et al., 2023). Surveying growth indicators across all individuals—both adults and juveniles—further allows assessment of how toxicity changes the growth distribution within a population, beyond comparisons of mean values. For example, Crouau and Moïa (2006) reported that the body-length distribution of *Folsomia candida* juveniles shifted from trimodal to unimodal with increasing cadmium concentration, and Marks et al. (2015) applied a multi-Gaussian model to distinguish cohorts from such distributions and to quantify their biomass, resolving structural changes undetectable by juvenile counts alone. Although body length has been the primary growth indicator, body area derived from image analysis has been reported across various invertebrate taxa to reflect actual biomass more accurately (Wardhaugh, 2013; Llopis-Belenguer et al., 2018). In *Caenorhabditis elegans*, body area and volume yielded lower EC_10_ values than body length, indicating higher sensitivity to toxicity (DuPlissis et al., 2025). Against this background, body area has begun to be explored as a growth indicator in Collembola toxicity tests (Guimarães et al., 2019a, 2019b, 2022).

Applying growth indicators, however, entails substantial methodological challenges. Collembola are very small, and juveniles in particular can number from hundreds to thousands per test unit, rendering individual measurement extremely time-consuming (Ding et al., 2021). Consequently, some studies have measured mean growth using only randomly selected juveniles (Marks et al., 2015; Hennig et al., 2022; Alves et al., 2023), an approach that does not represent the full size distribution of the population. Digital image-analysis software such as ImageJ has improved measurement precision, yet the labor and time required for individual imaging and manual measurement remain considerable (Bánszegi et al., 2014). Automation has been pursued separately for counting and for body-length measurement (Caridade et al., 2011; Bánszegi et al., 2014), but the two have rarely been combined within a single integrated workflow, so counting and growth assessment have remained separate. Wehrli et al. (2026) recently developed a YOLO-based automated counting program for Collembola. This program showed strong concordance with manual counts across diverse soil conditions in five laboratories across five countries. Furthermore, as a direction for advancing this methodology, they proposed the possibility of extending the approach to size-based classification and growth assessment, thereby realizing the simultaneous automation of counting and growth measurement within a single integrated workflow. To date, however, this integration has not been realized: automated counting has been validated, but the simultaneous, population-wide measurement of individual body size—and the population-structure information that such measurement carries—remains unavailable, leaving the richer and potentially more protective endpoints it would enable largely unexplored. This points to the need to move from the previously separate procedures of counting and growth evaluation toward an integrated automation framework, thereby suggesting a new trajectory for the advancement of AI-based ecotoxicological assessment.

In this study, we developed an AI-based system that couples YOLOv8 object detection with Voronoi–Otsu segmentation-based body-area measurement, so that counting and growth (body-area) assessment are performed automatically and simultaneously within a single workflow, and applied it to a metal(loid)-contaminated soil ecotoxicity test using *Allonychiurus kimi*, an Asian alternative indicator species. The objectives were to (1) validate the accuracy of the AI-based system for automated counting and body area measurement; (2) explore the utility of the resulting body area data as toxicity endpoints, including mean and cumulative body area; and (3) assess the effects of metal(loid)s on population structure through cohort analysis applying a multi-Gaussian model to the frequency distribution of juvenile body area. Through this approach, the study evaluates the applicability of an AI-based automated system to ecotoxicity testing and provides foundational data for understanding the responses of *A. kimi* to Cd, As, and Cu at the level of body area distribution. More broadly, it reframes the standard Collembola test from an abundance-only assay toward an integrated, population-structure–aware assessment whose more sensitive endpoints could translate into more protective effect concentrations for soil risk assessment, in a workflow that—like automated counting—is in principle transferable across species and laboratories.

## 2. Materials and Methods

### 2.1. Test organism

*A. kimi*, originally collected from a paddy field in Icheon, Gyeonggi Province, Korea, in 1996, has since been maintained in laboratory culture (accession PNUCOL001, Environmental Ecology Laboratory, Pusan National University, https://sites.google.com/view/ee-lab; Wee et al., 2021). Cultures were maintained in plastic Petri dishes (90 mm × 15 mm) on a moist substrate of plaster of Paris, activated charcoal, and distilled water (4:1:4, v/v/v) in continuous darkness at 20 ± 1 °C, and were fed weekly with Brewer’s yeast (Sigma-Aldrich, St. Louis, MO, USA). Age-synchronized adults were obtained by transferring eggs laid within a single week onto fresh substrate and rearing the hatchlings to the required age under identical conditions.

### 2.2. Toxicity test

The soil ecotoxicity test was conducted in accordance with OECD Test Guideline 232 (Collembola Reproduction Test in Soil; OECD, 2016) and ISO 11267 (Soil quality — Inhibition of Collembola Reproduction by Soil Contaminants; ISO, 2023), with modifications appropriate for evaluating As toxicity in *A. kimi* (Wee et al., 2021). Artificial soil was prepared on a dry-weight basis from sphagnum peat (< 2 mm; 5 %), kaolinite clay (20 %), and quartz sand (75 %), and the pH was adjusted to 6.0 ± 0.5 with CaCO_3_ (Sigma-Aldrich). Three metal(loid)s were tested: cadmium (Cd; CdCl_2_·2.5H_2_O) at 0, 12.5, 25, 50, 100, and 200 mg kg^−1^ (Son et al., 2011); arsenic (As; Na_2_HAsO_4_·7H_2_O) at 0, 10, 20, and 40 mg kg^−1^; and copper (Cu; CuCl_2_·2H_2_O) at 0, 50, 100, 200, 400, and 800 mg kg^−1^ (Son et al., 2017), all purchased from Sigma-Aldrich. Spiked soils were adjusted to 50 % of the water-holding capacity, dispensed (30 g) into PET vessels (160 mL), and aged for one week at 20 ± 1 °C. Ten 7–8-week-old adults were then introduced into each vessel, with five replicates per concentration. During the 28-day exposure, vessels were aerated weekly, moisture was replenished with distilled water, and approximately 10 mg of Brewer’s yeast was supplied biweekly.

After exposure, the soils were flooded with water and gently agitated to recover individuals by flotation; this was repeated at least six times until no further individuals surfaced. Adults (1.3–1.7 mm) and juveniles (< 0.8 mm) were distinguished by body length and counted manually under a stereomicroscope. The recovered individuals were then transferred onto charcoal substrates for image acquisition and AI-based analysis.

### 2.3. Image acquisition and dataset construction

The training and validation datasets to train and evaluate AI-based detection and body area measurement were independently constructed. In particular, the model was trained on individuals unrelated to the toxicity test and evaluated on individuals recovered from the actual test, thereby preventing data leakage and demonstrating applicability to realistic ecotoxicity testing conditions.

Training images were acquired independently of the toxicity test (Section 2.2) by inoculating *A. kimi* onto charcoal substrates across ten density conditions, ranging from 1 adult with 10 juveniles to 10 adults with 100 juveniles, with juveniles spanning four post-hatch ages (11, 17, 21, and 28 days); 40 images were captured at 5-minute intervals to form the initial training set. To improve juvenile detection, a supplementary set of 102 images was added, in which 50 and 100 juveniles were inoculated in triplicate and each plate was photographed 17 times at 5-minute intervals. The validation dataset was constructed from individuals recovered from the toxicity tests in Section 2.2: after manual counting, the adults and juveniles from each treatment were transferred onto separate charcoal substrates and photographed, yielding 160 images in total (6 Cd, 4 As, and 6 Cu concentrations, each with five replicates, with adults and juveniles imaged separately).

All imaging was conducted on plastic Petri dishes (90 mm in diameter) filled with a plaster of Paris–charcoal–water substrate (4:1:4, v/v/v), with a 5 mm × 5 mm white marker placed on each substrate for pixel-to-area conversion. Images were captured under standardized distance and lighting conditions using a Canon EOS M digital camera (Canon Inc., Tokyo, Japan). The captured images were preprocessed in Adobe Photoshop (Adobe Inc., San Jose, CA, USA) to remove the background and non-target debris, and edge masking was applied to the substrate boundary. Roboflow (https://roboflow.com/), an online labeling tool by drawing a bounding box for an object and assign a user-defined class, was employed to label the adults and juveniles in the images and to obtain the structured dataset to be used as the inputs of the YOLOv8 model for detecting adults and juveniles. The whole labeling process was conducted by a single annotator to ensure consistency.

### 2.4. AI-based detection and body area measurement

Object detection was performed using the YOLOv8 model (Varghese and Sambath, 2024). Model evaluation was conducted at two distinct levels: the object-detection performance of the model (this section), and the agreement between the resulting individual counts and body area measurements and the corresponding manual values (Section 2.5). The performance of detection model for adults and juveniles is measured by five popular metrics: precision, recall, mean average precision at an Intersection over Union (IoU) threshold of 0.5 (mAP50), mean average precision across IoU thresholds from 0.5 to 0.95 (mAP50-95), and the *F*1-score. A true positive (TP) was defined as a predicted bounding box overlapping the ground-truth box with an IoU of at least 0.666 and assigned to the correct class. Precision, recall, and the *F*1-score were calculated using Eqs. 1, 2, and 5, and the mean average precision metrics (mAP50 and mAP50-95) using Eqs. 3 and 4.

(Eq. 1)

where FP denotes the number of false positives.

(Eq. 2)

where FN denotes the number of false negatives (existing individuals not detected or misclassified).

(Eq. 3)

where C is the number of classes and AP the average precision (the area under the precision–recall curve) for class c, evaluated at an IoU threshold of 0.5.

(Eq. 4)

where N is the number of IoU thresholds (t = 0.5, 0.55, …, 0.95) and AP the average precision for class c at threshold t.

(Eq. 5)

All bounding boxes for the detected adults and juveniles in an image are cropped and each cropped image is fed into the pipeline of body area estimation, which consists of five steps: (i) converting the cropped image to grayscale, (ii) binarization (i.e., thresholding each pixel into either 0 (black) or 1 (white) based on a predefined grayscale value), (iii) applying Voronoi–Otsu segmentation to identify the segments in the given image, (iv) choosing the segment closest to the center of the image, and (v) computing the body area of the detected object by multiplying the scaling factor to the number of the pixels of the detected object. Here, the scaling factor is calculated on the number of pixels of the 5 mm × 5 mm marker (0.0025 mm^2^ pixel^−1^). We conducted a grid-search method to find the best hyperparameter set for the detection and segmentation in different test substances (i.e., Cd, As, and Cu) and object type (i.e., adult and juvenile). The selected hyperparameter sets are described in Table 1. Representative validation images and the corresponding detection results are shown in Fig. S1.

**Table 1.**
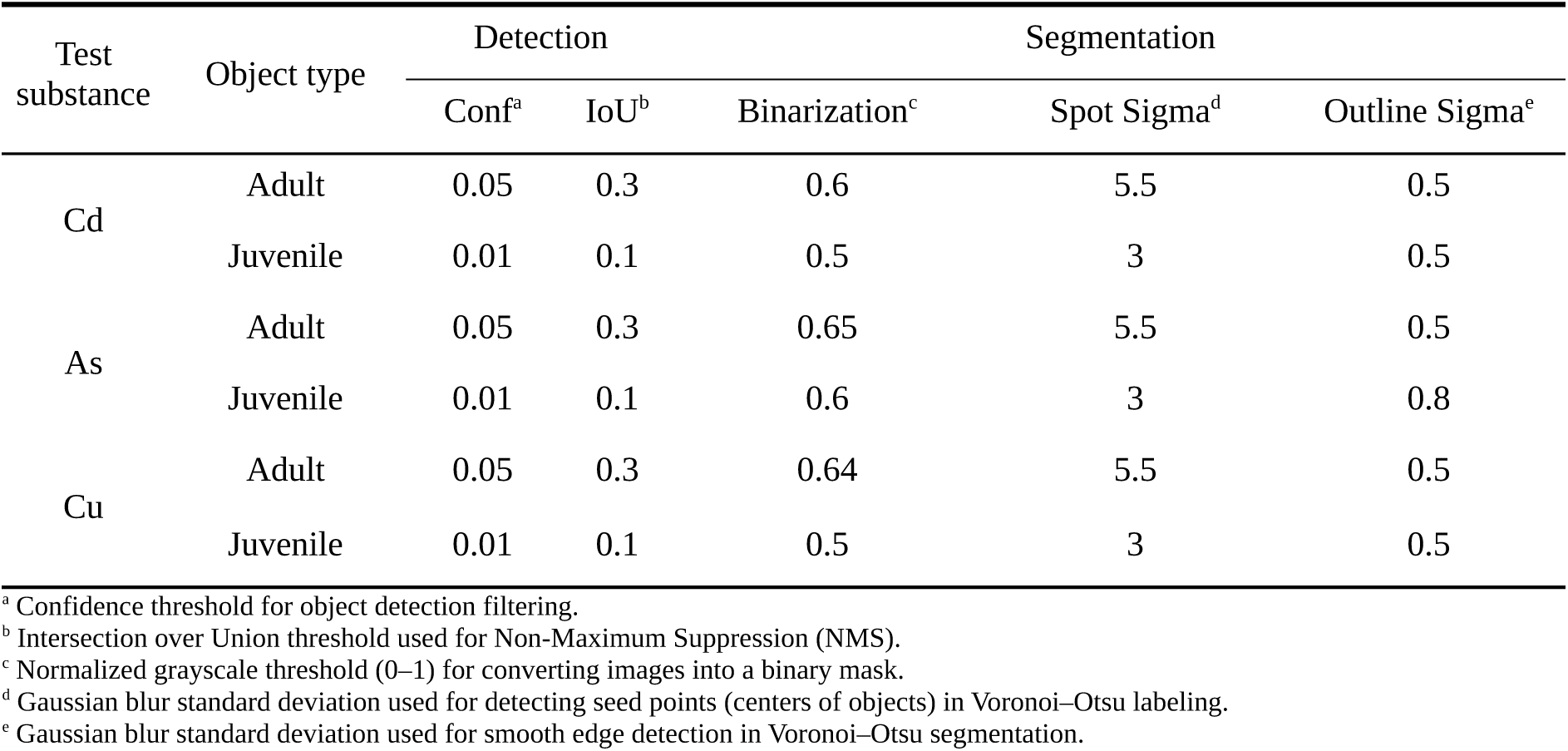
The selected hyperparameter sets of detection and segmentation for different test substances and object types.

### 2.5. Agreement between manual and AI-based measurements

To assess the objective performance of the model, the accuracy analysis in this section used the full set of AI-detected individuals without filtering; for counting, the total number of individuals detected by the model—irrespective of bounding-box localization quality—was compared with the corresponding manual count for each sample. Manual counting was performed under a stereomicroscope as described in Section 2.2, and manual measurements of body area, body length, and body width were made on AI-cropped single-individual images using ImageJ (NIH, Bethesda, MD, USA). Each image was converted to 8-bit format and binarized using a treatment-specific threshold, followed by visual inspection and manual adjustment to ensure that the segmented region accurately matched the body outline of each individual. To compare body area with the other growth indicators, the *Analyze Particles* function, with the *Fit ellipse* option enabled, was then used to measure the body area, major axis (body length), and minor axis (body width) of each individual.

Agreement between the two methods was evaluated using the coefficient of determination (*R*^2^) and the relative root mean square error (*r*RMSE). With the manual values treated as the reference and the AI-based values as the predictions, *R*^2^ was computed in the model-efficiency form (one minus the ratio of the residual sum of squares to the total sum of squares), which expresses agreement relative to the 1:1 identity line (Nash and Sutcliffe, 1970). The *r*RMSE was expressed as the relative error with respect to the manual mean, and model performance was classified as Excellent (< 10 %), Good (10–20 %), Weak (20–30 %), or Poor (> 30 %) following Despotovic et al. (2016). These two metrics were calculated using Eqs. 6 and 7, respectively.

(Eq. 6)

where M and A are the paired manual and AI-based values and M^Y^ the mean of the manual values; a value of 1 indicates perfect agreement with the 1:1 line.

(Eq. 7)

where n is the number of paired values; *r*RMSE is expressed as a percentage of the manual mean.

### 2.6. Cohort structure analysis of juvenile body area distribution

The cohort analysis and the downstream endpoint analyses (Section 2.7) were performed on AI-measured juvenile data after excluding detections with unambiguous overestimation of body area, such as overlapping individuals segmented as a single object. The detections were identified by visual inspection of each cropped image, independently of the manual reference measurements, and accounted for 0.58 % (As), 0.84 % (Cd), and 2.39 % (Cu) of juvenile detections; no merged detections occurred among adults.

For each concentration, a Gaussian mixture model (GMM) was fitted independently to the mm^2^-converted body area distribution (Section 2.4) using the mclust package in R version 4.3.1 (R Core Team, 2023; Fraley et al., 2012). One to six Gaussian components were considered as candidates, each allowed its own variance (the “V” model), and the optimal model was selected by the Bayesian Information Criterion (BIC). The mean and standard deviation of each component were extracted from the selected model, and cohort boundaries were defined as the midpoints between adjacent component means. Because all concentrations yielded no more than two components, individuals were assigned to cohorts (Cohort 1, smaller body area; Cohort 2, larger body area).

To verify the validity of the cohort separation, any concentration resolved into more than one component was evaluated against the model with one fewer component. Specifically, the difference in BIC relative to the single-component model (G = 1; ΔBIC) was calculated, and a sequential bootstrap likelihood-ratio test (mclustBootstrapLRT, “V” model, 9999 bootstrap replications; McLachlan, 1987; McLachlan and Rathnayake, 2014) was conducted to test whether each additional component yielded a significant improvement in fit.

### 2.7. Toxicity endpoints

To compare the sensitivity of different toxicity endpoints, three endpoints were derived for each treatment: the number of individuals, the average body area, and the cumulative body area. The cumulative body area was defined as the sum of the body areas of all individuals in a given replication, providing an integrative endpoint that simultaneously reflects abundance and individual size. The average body area was calculated as the mean of the individual body areas (in mm^2^). Each endpoint was calculated separately for adults and juveniles. For the number of individuals, the adult and juvenile values correspond to the conventional endpoints of the Collembola reproduction test—adult survival and reproduction (juvenile production), respectively.

### 2.8. Statistical analysis

All statistical analyses were performed in R version 4.3.1 (R Core Team, 2023) and SAS version 9.4 (SAS Institute Inc., Cary, NC, USA), with statistical significance accepted at *p* < 0.05. The agreement between the manual and AI-based measurements was assessed using *R*^2^ and *r*RMSE as defined in Section 2.5, and the cohort structure of the juvenile body-area distribution was resolved by Gaussian mixture modeling as described in Section 2.6. The concentration–response analysis of the toxicity endpoints is described below.

Concentration effects on each endpoint were evaluated by one-way analysis of variance (ANOVA) followed by Dunnett’s test against the control, from which the no-observed-effect concentration (NOEC) and the lowest-observed-effect concentration (LOEC) were determined. EC_50_ values for each endpoint were estimated by fitting the Haanstra logistic dose–response model (Haanstra et al., 1985; Eq. 8) with the minpack.lm package in R, and are reported with their 95 % confidence intervals (CIs). Differences in sensitivity among the three endpoints were tested by comparing their EC_50_ estimates (Wald Z-test; Eq. 9).

(Eq. 8)

where y is the response, x = log_10_(concentration + 1), c the control-level response, b the slope, and a the log-scale midpoint (corresponding to EC_50_).

(Eq. 9)

where a_1_ and a_2_ are the log-scale midpoints of the two endpoints and SE_1_ and SE_2_ their standard errors; the test is two-sided, with *p* < 0.05 considered significant.

## 3. Results

### 3.1. Performance of the AI-based automated measurement system

Table 2 summarizes the detection performance of the AI-based model for adults and juveniles, including precision, recall, mAP50, mAP50-95, and the *F*1-score. For adults, a total of 601 instances were detected from 600 actual individuals across 76 images, with one false positive. The model achieved a precision of 1.000, recall of 0.950, mAP50 of 0.990, mAP50-95 of 0.672, and an *F*1-score of 0.974. For juveniles, 3,033 instances were detected from 3,124 actual individuals across 69 images, with a precision of 0.950, recall of 0.830, mAP50 of 0.929, mAP50-95 of 0.523, and an *F*1-score of 0.886. Overall, both adults and juveniles exhibited high precision and mAP50 values. Adults in particular showed near-perfect detection performance, with a precision of 1.000. Although juveniles had lower recall than adults, their detection accuracy remained high, as reflected by their high precision.

**Table 2.**
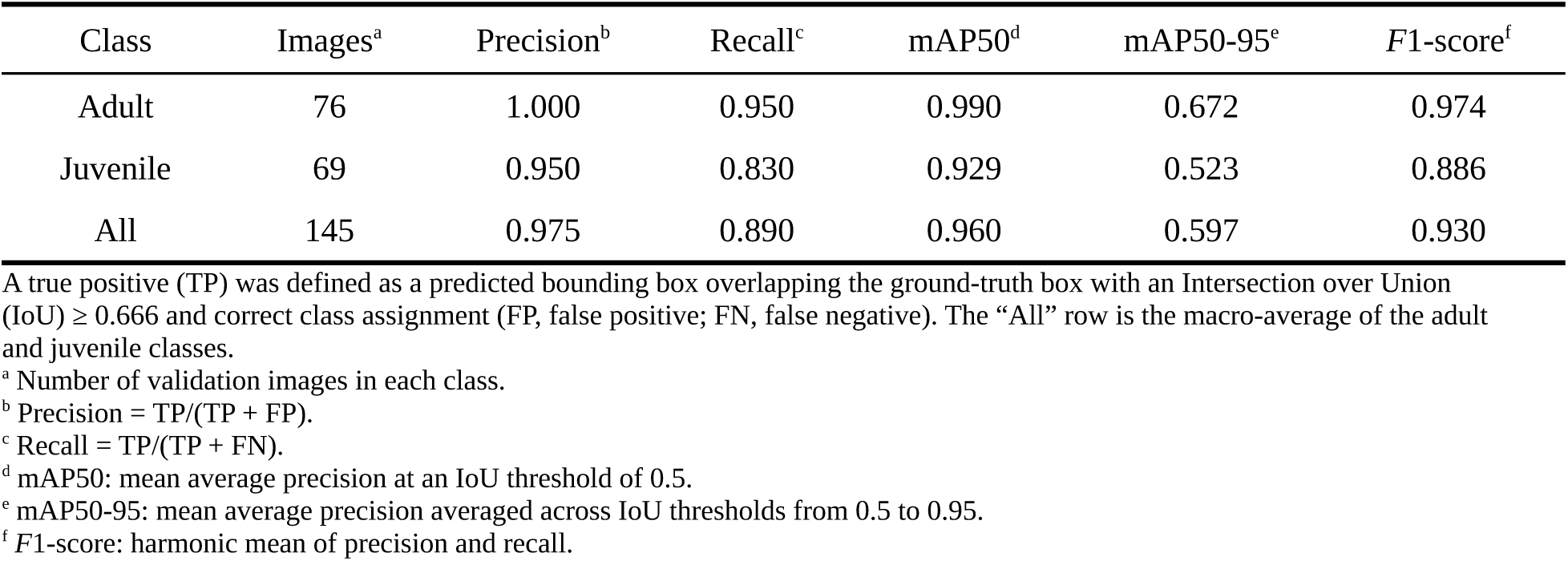
Object-detection performance of the AI model for adult and juvenile Collembola.

| Class | Images<sup>a</sup> | Precision<sup>b</sup> | Recall<sup>c</sup> | mAP50<sup>d</sup> | mAP50-95<sup>e</sup> | F1-score<sup>f</sup> |
| --- | --- | --- | --- | --- | --- | --- |
| Adult | 76 | 1.000 | 0.950 | 0.990 | 0.672 | 0.974 |
| Juvenile | 69 | 0.950 | 0.830 | 0.929 | 0.523 | 0.886 |
| All | 145 | 0.975 | 0.890 | 0.960 | 0.597 | 0.930 |

A true positive (TP) was defined as a predicted bounding box overlapping the ground-truth box with an Intersection over Union (IoU) $\geq 0.666$ and correct class assignment (FP, false positive; FN, false negative). The “All” row is the macro-average of the adult and juvenile classes.
<sup>a</sup> Number of validation images in each class.
<sup>b</sup> Precision = TP/(TP + FP).
<sup>c</sup> Recall = TP/(TP + FN).
<sup>d</sup> mAP50: mean average precision at an IoU threshold of 0.5.
<sup>e</sup> mAP50-95: mean average precision averaged across IoU thresholds from 0.5 to 0.95.
<sup>f</sup> F1-score: harmonic mean of precision and recall.

Building on this detection performance, the agreement between AI-based automated measurements and manual measurements was evaluated for each metal(loid) by integrating the concentration gradient for the three toxic metal(loid)s (Cd, As, and Cu) (Table 3). Manual and automated mean values were calculated as abundance-weighted averages across all concentrations, and the agreement between the two methods was quantified using *r*RMSE and *R*^2^. For individual counting, all metal(loid)s and developmental stages showed very high agreement, with *R*^2^ values ranging from 0.99 to 1.00. In adults, the *r*RMSE was 3.18 % for Cd and 0.00 % for both As and Cu. In juveniles, the *r*RMSE values were 5.49 % for Cd, 8.87 % for As, and 3.09 % for Cu. For body area measurement, all metal(loid)s and developmental stages showed high agreement, with *R*^2^ values ranging from 0.95 to 1.00. In adults, the *r*RMSE values were 2.18 % for Cd, 1.74 % for As, and 1.60 % for Cu. In juveniles, they were 7.39 % for Cd, 9.52 % for As, and 6.58 % for Cu. All *r*RMSE values for both individual counting and body area measurement were below 10 %, placing them in the “Excellent” category according to the model performance criteria of Despotovic et al. (2016). In adults, agreement was particularly high, with *r*RMSE values at or below 3.18 % across all endpoints.

**Table 3.** Agreement between manual and AI-based automated measurements of individual counts and body area for *A. kimi* exposed to cadmium (Cd), arsenic (As), and copper (Cu).

| Indicating value | Test substance | Developmental Stage | $r\text{RMSE}^a$ | $R^{2b}$ | Manual Mean ± SD<sup>c</sup> | Automatic Mean ± SD |
| --- | --- | --- | --- | --- | --- | --- |
| Count | Cd | Adult | 3.18 | 1.00 | 5.73 ± 3.37 | 5.77 ± 3.4 |
| | | Juvenile | 5.49 | 1.00 | 28.8 ± 27.83 | 28.03 ± 26.82 |
| | As | Adult | 0.00 | 1.00 | 7.25 ± 3.92 | 7.25 ± 3.92 |
| | | Juvenile | 8.87 | 0.99 | 27.5 ± 22.01 | 25.65 ± 20.84 |
| | Cu | Adult | 0.00 | 1.00 | 9.43 ± 0.84 | 9.43 ± 0.84 |
| | | Juvenile | 3.09 | 1.00 | 57 ± 28.88 | 55.97 ± 28.27 |
| Body area (mm<sup>2</sup>) | Cd | Adult | 2.18 | 1.00 | 0.497 ± 0.197 | 0.498 ± 0.192 |
| | | Juvenile | 7.39 | 0.98 | 0.089 ± 0.042 | 0.091 ± 0.042 |
| | As | Adult | 1.74 | 0.99 | 0.550 ± 0.134 | 0.556 ± 0.135 |
| | | Juvenile | 9.52 | 0.95 | 0.083 ± 0.026 | 0.086 ± 0.023 |
| | Cu | Adult | 1.60 | 0.99 | 0.696 ± 0.156 | 0.690 ± 0.152 |
| | | Juvenile | 6.58 | 0.97 | 0.182 ± 0.073 | 0.184 ± 0.071 |

Values are abundance-weighted means ( ± SD) integrated across all test concentrations for each metal(loid) and developmental stage. Counts are expressed as the number of individuals, and body area in mm<sup>2</sup>.
<sup>a</sup> Relative root mean square error.
<sup>b</sup> Coefficient of determination relative to the 1:1 line.
<sup>c</sup> Standard deviation.

On a per-concentration basis, agreement for body area fell within the “Excellent” category (*r*RMSE < 10 %) for all treatments except two low-abundance juvenile treatments—25 mg kg^−1^ Cd (15.09 %) and 10 mg kg^−1^ As (11.66 %)—which were classified as “Good” (*r*RMSE 10–20 %; Tables S1–S3).

### 3.2. Cohort structure analysis of juvenile body area distribution

Across all treatments, the juvenile body area distribution was resolved by GMM into at most two cohorts—Cohort 1 (smaller individuals) and Cohort 2 (larger individuals) (Fig. 1; Table 4). In every treatment resolved into two cohorts, the two-cohort model was significantly better supported than the single-cohort model (ΔBIC > 2, indicating at least positive evidence sensu Kass and Raftery (1995); bootstrap likelihood-ratio test, *p* < 0.001; Table S4), confirming that the cohort separation was not an artifact of over-splitting. For all three metal(loid)s, the proportion of Cohort 2 progressively decreased with increasing concentration, becoming minimal or disappearing entirely—leaving a single-cohort structure composed only of Cohort 1—at the highest concentrations.

**Fig. 1.**
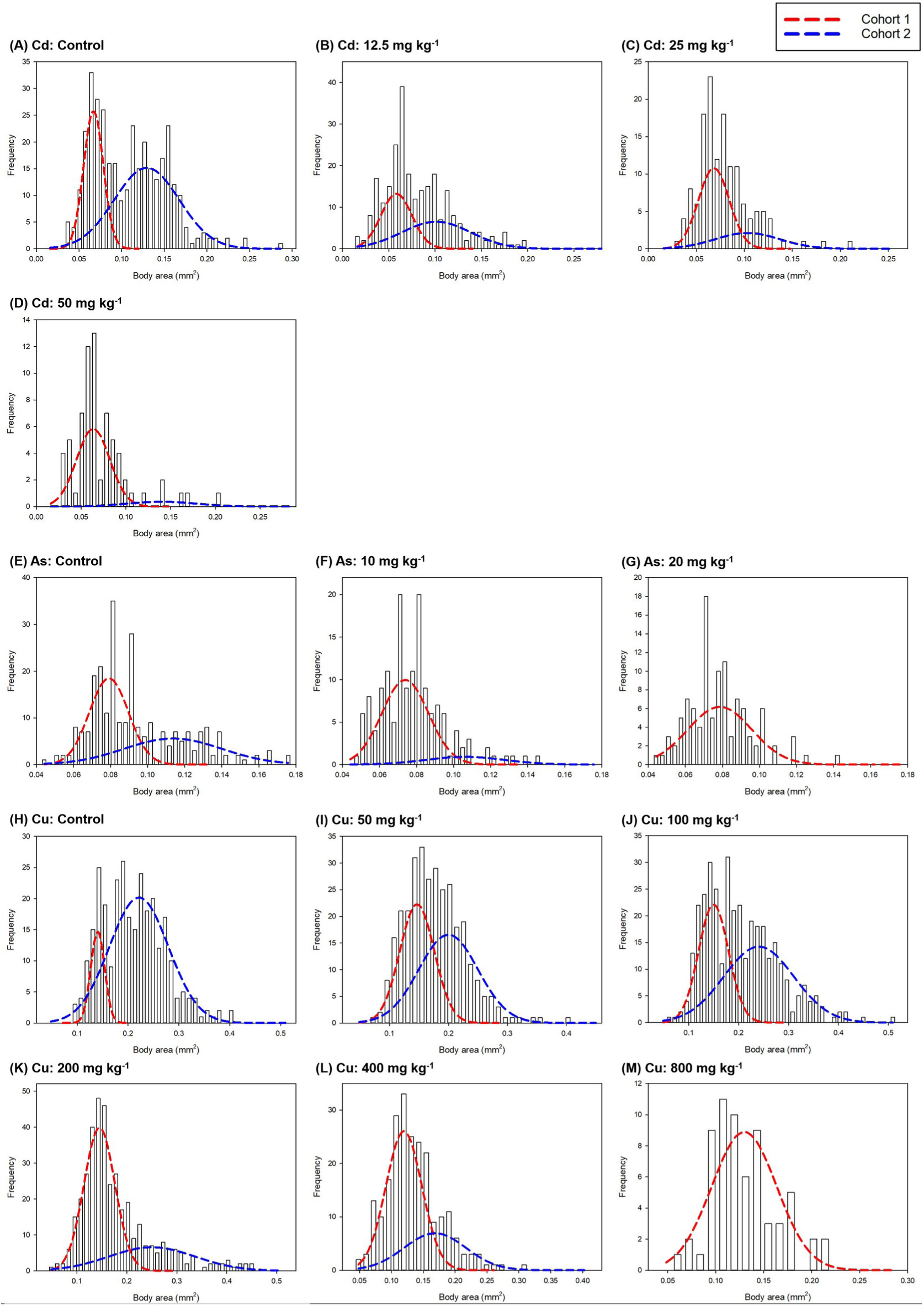
Body area frequency distributions of *Allonychiurus kimi* juveniles across metal(loid) concentrations, decomposed into two cohorts by Gaussian mixture modeling (GMM). Panels show cadmium (Cd; A–D), arsenic (As; E–G), and copper (Cu; H–M) treatments. Bars indicate observed frequencies, and dashed curves represent the fitted GMM components (red, Cohort 1 [smaller body area]; blue, Cohort 2 [larger body area]). GMM was not fitted where too few juveniles were recovered for reliable decomposition (Cd, 100 and 200 mg kg^−1^); no juveniles were recovered at 40 mg kg^−1^ As.

**Table 4.**
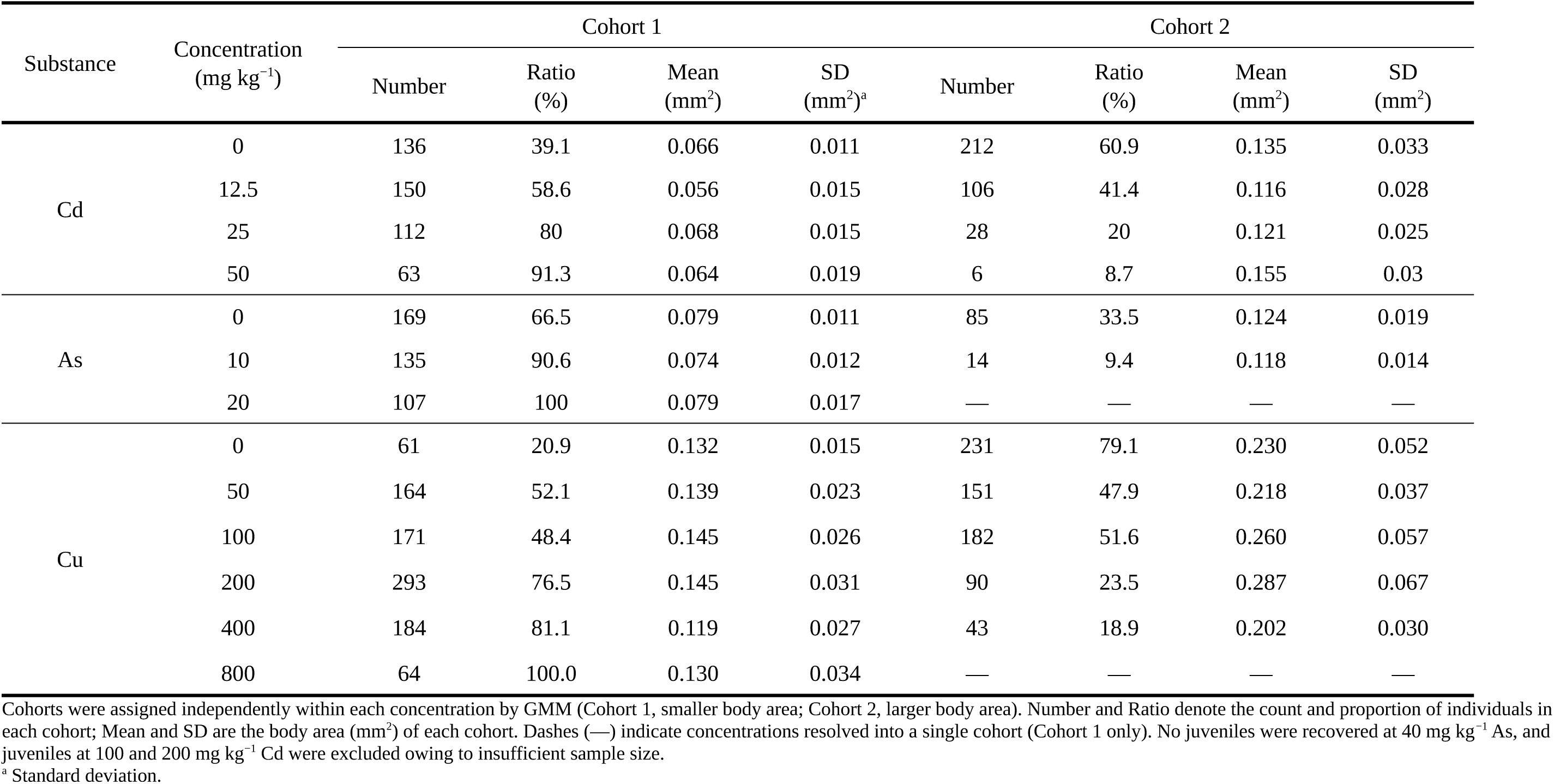
Cohort structure of juvenile *A. kimi* body area resolved by Gaussian mixture modeling (GMM) following exposure to cadmium (Cd), arsenic (As), and copper (Cu).

| Substance | Concentration<br>(mg kg<sup>-1</sup>) | Cohort 1 |  |  |  | Cohort 2 |  |  |  |
| --- | --- | --- | --- | --- | --- | --- | --- | --- | --- |
|  |  | Number | Ratio<br>(%) | Mean<br>(mm<sup>2</sup>) | SD<br>(mm<sup>2</sup>)<sup>a</sup> | Number | Ratio<br>(%) | Mean<br>(mm<sup>2</sup>) | SD<br>(mm<sup>2</sup>) |
| Cd | 0 | 136 | 39.1 | 0.066 | 0.011 | 212 | 60.9 | 0.135 | 0.033 |
|  | 12.5 | 150 | 58.6 | 0.056 | 0.015 | 106 | 41.4 | 0.116 | 0.028 |
|  | 25 | 112 | 80 | 0.068 | 0.015 | 28 | 20 | 0.121 | 0.025 |
|  | 50 | 63 | 91.3 | 0.064 | 0.019 | 6 | 8.7 | 0.155 | 0.03 |
| As | 0 | 169 | 66.5 | 0.079 | 0.011 | 85 | 33.5 | 0.124 | 0.019 |
|  | 10 | 135 | 90.6 | 0.074 | 0.012 | 14 | 9.4 | 0.118 | 0.014 |
|  | 20 | 107 | 100 | 0.079 | 0.017 | — | — | — | — |
| Cu | 0 | 61 | 20.9 | 0.132 | 0.015 | 231 | 79.1 | 0.230 | 0.052 |
|  | 50 | 164 | 52.1 | 0.139 | 0.023 | 151 | 47.9 | 0.218 | 0.037 |
|  | 100 | 171 | 48.4 | 0.145 | 0.026 | 182 | 51.6 | 0.260 | 0.057 |
|  | 200 | 293 | 76.5 | 0.145 | 0.031 | 90 | 23.5 | 0.287 | 0.067 |
|  | 400 | 184 | 81.1 | 0.119 | 0.027 | 43 | 18.9 | 0.202 | 0.030 |
|  | 800 | 64 | 100.0 | 0.130 | 0.034 | — | — | — | — |

Cohorts were assigned independently within each concentration by GMM (Cohort 1, smaller body area; Cohort 2, larger body area). Number and Ratio denote the count and proportion of individuals in each cohort; Mean and SD are the body area (mm<sup>2</sup>) of each cohort. Dashes (—) indicate concentrations resolved into a single cohort (Cohort 1 only). No juveniles were recovered at 40 mg kg<sup>-1</sup> As, and juveniles at 100 and 200 mg kg<sup>-1</sup> Cd were excluded owing to insufficient sample size.
<sup>a</sup> Standard deviation.

Following As exposure, the proportion of Cohort 2 declined sharply from 33.5 % in the control (85 individuals, 0.124 ± 0.019 mm^2^) to 9.4 % at 10 mg kg^−1^ (14 individuals, 0.118 ± 0.014 mm^2^) and was no longer observed at 20 mg kg^−1^. Correspondingly, the proportion of Cohort 1 increased from 66.5 % in the control to 90.6 % at 10 mg kg^−1^ and 100 % at 20 mg kg^−1^, while its mean body area remained within 0.074–0.079 mm^2^. The number of Cohort 1 individuals decreased with increasing concentration, from 169 in the control to 135 at 10 mg kg^−1^ and 107 at 20 mg kg^−1^.

For Cd, the proportion of Cohort 2 likewise decreased continuously with increasing concentration, from 60.9 % in the control (212 individuals, 0.135 ± 0.033 mm^2^) to 41.4 % at 12.5 mg kg^−1^ (106 individuals, 0.116 ± 0.028 mm^2^), 20.0 % at 25 mg kg^−1^ (28 individuals, 0.121 ± 0.025 mm^2^), and 8.7 % at 50 mg kg^−1^ (6 individuals, 0.155 ± 0.030 mm^2^). The proportion of Cohort 1 correspondingly increased from 39.1 % in the control to 91.3 % at 50 mg kg^−1^, with a mean body area of 0.056–0.068 mm^2^. The number of Cohort 1 individuals first increased from 136 in the control to 150 at 12.5 mg kg^−1^ and then decreased to 112 at 25 mg kg^−1^ and 63 at 50 mg kg^−1^.

For Cu, the proportion of Cohort 2 decreased from 79.1 % in the control (231 individuals, 0.230 ± 0.052 mm^2^) to 47.9 % at 50 mg kg^−1^ (151 individuals, 0.218 ± 0.037 mm^2^) and 51.6 % at 100 mg kg^−1^ (182 individuals, 0.260 ± 0.057 mm^2^), then declined further to 23.5 % at 200 mg kg^−1^ (90 individuals, 0.287 ± 0.067 mm^2^) and 18.9 % at 400 mg kg^−1^ (43 individuals, 0.202 ± 0.030 mm^2^), and was no longer observed at 800 mg kg^−1^. The proportion of Cohort 1 increased from 20.9 % in the control to 100 % at 800 mg kg^−1^. In contrast to both cohorts for As and Cd and to Cohort 2 for Cu—all of which decreased in number with increasing concentration—the number of Cohort 1 individuals exposed to Cu increased from 61 in the control to 164 at 50 mg kg^−1^, 171 at 100 mg kg^−1^, and 293 at 200 mg kg^−1^ (an approximately 4.8-fold increase) before decreasing to 184 at 400 mg kg^−1^ and 64 at 800 mg kg^−1^.

### 3.3. Comparison of toxicity assessment based on the number of individuals, average body area, and cumulative body area

Across all three metal(loid)s, the number of individuals, average body area, and cumulative body area all decreased with increasing concentration, but the rate of decline and the concentration at which significant effects first appeared differed among the three endpoints (Fig. 2). The concentration– response patterns are described below for each metal(loid).

**Fig. 2.**
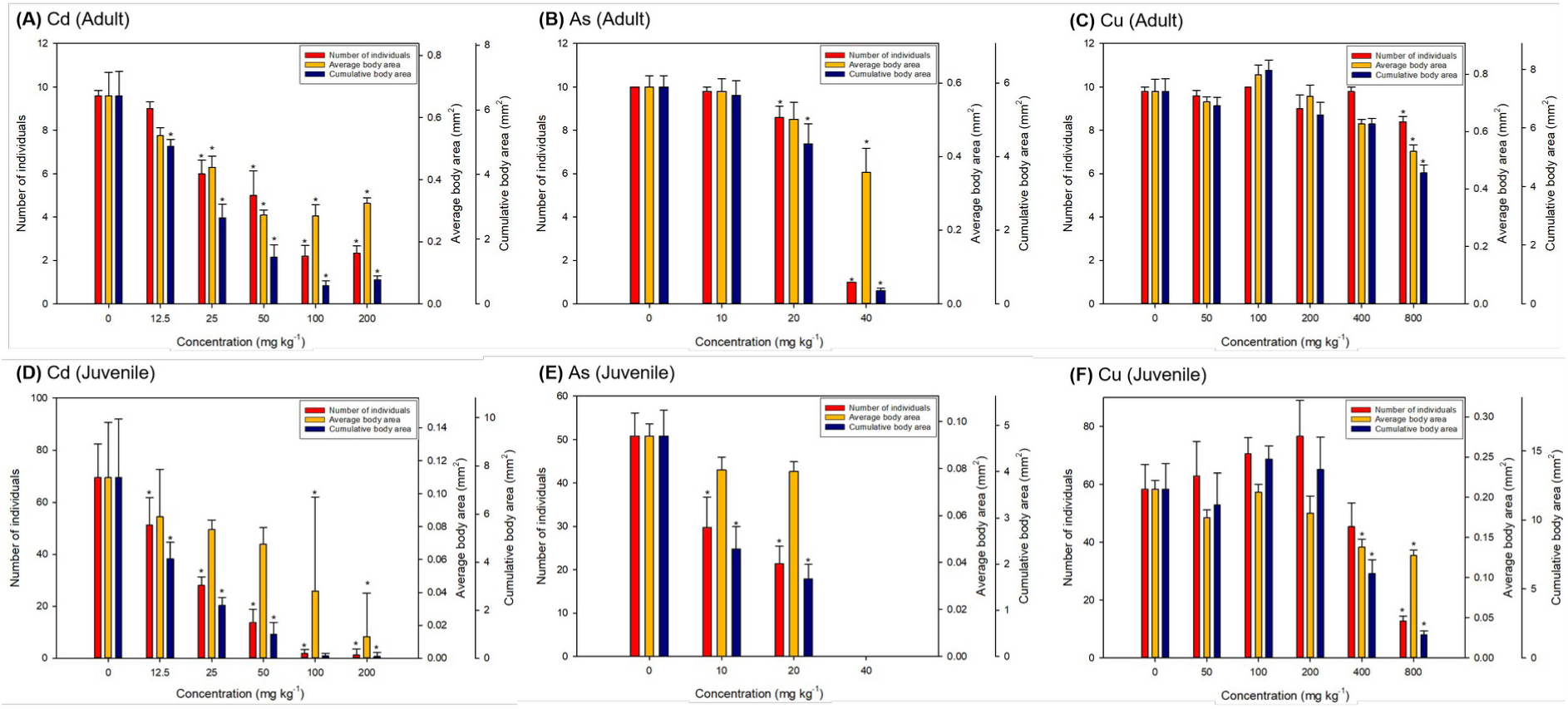
Concentration–response of three toxicity endpoints in *Allonychiurus kimi* exposed to cadmium (Cd), arsenic (As), and copper (Cu): number of individuals (red; left axis), average body area (yellow), and cumulative body area (blue) (right axes). Panels show adults (A–C) and juveniles (D–F). Bars represent treatment means and error bars the standard deviation; asterisks (*) denote significant differences from the control (Dunnett’s test, *p* < 0.05).

For Cd, the number of individuals, average body area, and cumulative body area of adults were 9.60 individuals, 0.67 ± 0.15 mm^2^, and 6.44 ± 1.51 mm^2^ in the control, decreasing to 1.40 individuals, 0.19 ± 0.16 mm^2^, and 0.45 ± 0.39 mm^2^ at the highest concentration (200 mg kg^−1^) (Fig. 2A). Cumulative body area was the most sensitive endpoint, with its LOEC (12.5 mg kg^−1^) one concentration step below that of the number of individuals and average body area (both 25 mg kg ^−1^). In juveniles, the corresponding values declined from 69.60 individuals, 0.11 ± 0.03 mm^2^, and 7.53 ± 2.41 mm^2^ in the control to 1.20 individuals, 0.01 ± 0.03 mm^2^, and 0.08 ± 0.16 mm^2^ at 200 mg kg^−1^ (Fig. 2D). The number of individuals and cumulative body area shared the same LOEC (12.5 mg kg^−1^), whereas that of average body area was three concentration steps higher (100 mg kg^−1^).

For As, the three endpoints in adults were 10.00 individuals, 0.59 ± 0.06 mm^2^, and 5.89 ± 0.61 mm^2^ in the control, decreasing to 0.60 individuals, 0.22 ± 0.19 mm^2^, and 0.22 ± 0.19 mm^2^ at 40 mg kg^−1^ (Fig. 2B). The number of individuals and cumulative body area shared the same LOEC (20 mg kg ^−1^), whereas average body area was one step less sensitive (40 mg kg^−1^). In juveniles, the corresponding values changed from 50.80 individuals, 0.09 ± 0.01 mm^2^, and 4.77 ± 1.12 mm^2^ in the control to 21.40 individuals, 0.08 ± 0.01 mm^2^, and 1.68 ± 0.62 mm^2^ at 20 mg kg^−1^, with no individuals observed at 40 mg kg^−1^ (Fig. 2E). The number of individuals and cumulative body area shared the same LOEC (10 mg kg^−1^), whereas average body area showed no significant difference from the control at any concentration up to 20 mg kg^−1^, the highest concentration at which juveniles were recovered.

For Cu, the three endpoints in adults were 9.80 individuals, 0.74 ± 0.08 mm^2^, and 7.25 ± 0.87 mm^2^ in the control, decreasing to 8.40 individuals, 0.53 ± 0.05 mm^2^, and 4.47 ± 0.55 mm^2^ at 800 mg kg^−1^, with all three endpoints sharing the same LOEC (800 mg kg^−1^) (Fig. 2C). In juveniles, the corresponding values declined from 58.40 individuals, 0.21 ± 0.02 mm^2^, and 12.24 ± 3.71 mm^2^ in the control to 12.80 individuals, 0.13 ± 0.01 mm^2^, and 1.66 ± 0.57 mm^2^ at 800 mg kg^−1^ (Fig. 2F). The number of individuals and cumulative body area increased up to 200 mg kg^−1^ before declining, although the increase was not statistically significant (*p* > 0.05), whereas average body area decreased gradually without an increasing phase. Average body area and cumulative body area shared the same LOEC (400 mg kg^−1^), one step more sensitive than that of the number of individuals (800 mg kg^−1^).

A comparison of the EC_50_ values derived from the three juvenile endpoints (Table 5) showed that, for all metal(loid)s, cumulative body area yielded the lowest EC_50_ and average body area the highest. The EC_50_ values for cumulative body area, the number of individuals, and average body area in juveniles were 14.10, 21.09, and 65.28 mg kg^−1^ for Cd; 10.53, 13.60, and 23.87 mg kg^−1^ for As; and 398.80, 513.72, and 1223.36 mg kg^−1^ for Cu, respectively. A Z-test confirmed significant differences for Cd between the number of individuals and average body area (*p* = 0.006) and between average body area and cumulative body area (*p* = 0.001), with the difference between the number of individuals and cumulative body area approaching significance (*p* = 0.051). Significant differences were similarly observed for Cu (*p* = 0.026 and 0.003 for the corresponding pairs), whereas no significant differences among the endpoints were found for As (*p* = 0.435–0.586).

**Table 5.** EC_50_ values with 95 % confidence intervals for three toxicity endpoints in juvenile *Allonychiurus kimi* exposed to cadmium (Cd), arsenic (As), and copper (Cu).

| Test substance | Endpoint | EC<sub>50</sub> (mg kg<sup>-1</sup>) | 95 % CI lower | 95 % CI upper |
| --- | --- | --- | --- | --- |
| Cd | Number of individuals | 21.09 | 17.21 | 25.81 |
|  | Average body area | 65.28 | 29.86 | 141.35 |
|  | Cumulative body area | 14.10 | 9.86 | 19.99 |
| As | Number of individuals | 13.60 | 8.95 | 20.43 |
|  | Average body area | 23.87 | 2.80 | 161.96 |
|  | Cumulative body area | 10.53 | 6.33 | 17.13 |
| Cu | Number of individuals | 513.72 | 379.29 | 695.67 |
|  | Average body area | 1223.36 | 608.33 | 2459.14 |
|  | Cumulative body area | 398.80 | 323.28 | 491.90 |

EC<sub>50</sub> and 95 % confidence interval (CI) bounds are expressed in mg kg<sup>-1</sup>, estimated using the Haanstra model with concentration transformed as log<sub>10</sub>(concentration + 1). “Number of individuals” denotes the number of juveniles.

Across the three metal(loid)s, cumulative body area was consistently the most sensitive endpoint, showing the lowest EC_50_ among all endpoints (Table S5). In a parallel EC_50_ comparison based on manual measurements, body area was as sensitive as, or more sensitive than, body length and body width depending on the metal(loid), and the EC_50_ estimates for average body area showed more stable confidence intervals than those for body length and body width.

## 4. Discussion

### 4.1. Performance of the AI-based model

The AI-based object detection model developed in this study detected *A. kimi* individuals with high precision and recall across all developmental stages and metal(loid) exposure conditions. Detection performance was nonetheless lower for juveniles than for adults, which likely stems from the inherent limitations of small-object detection. Because small targets provide limited pixel information, the features available for classification and localization within YOLO-type convolutional neural network architectures are reduced (Vasanthi and Mohan, 2024; Nguyen et al., 2020). Moreover, the reproduction test produces juveniles in large numbers (Ding et al., 2021), so that individuals are frequently densely clustered and overlapping on the confined substrate. When many such small individuals overlap (occlusion), separating them into distinct objects becomes difficult and false negatives tend to increase (Nikouei et al., 2025; Aldubaikhi and Patel, 2025; Kos et al., 2023). Architectural improvements—such as multi-scale feature extraction and attention-based modules—are therefore needed to enhance small-object detection in future work.

The high agreement between manual and AI-based automated counts indicates that the approach can reliably reproduce the standard toxicity endpoints required by the OECD 232 reproduction test, and the comparably high agreement for body area across all metal(loid) concentrations confirms that the model performs stably across diverse environmental conditions and stressors. Previous work has shown that YOLO-based automated counting markedly reduces analysis time—by approximately 97 % relative to manual counting for *Folsomia candida* (Wehrli et al., 2026)—while achieving comparable accuracy, and automated counting approaches more broadly improve analytical efficiency (Marçal and Caridade, 2006; Caridade et al., 2011; Oriol et al., 2024). The present results reconfirm this validity in soil invertebrate ecotoxicity assessment. By minimizing variability arising from observer fatigue and subjective judgment, the automated approach improves data consistency and reproducibility (Sys et al., 2022), and by enabling simultaneous counting and body area measurement across many samples, it demonstrates clear potential for high-throughput ecotoxicity assessment.

In AI-based automation for Collembola, individual counting and growth measurement have developed as separate research streams, and few systems have integrated both functions. COLLEMBOT (Wehrli et al., 2026) marked an important advance by establishing a sophisticated automated counting system for *Folsomia candida* and validating its potential integration into the OECD 232 workflow, while outlining a direction in which growth-related information could be obtained through body area estimation based on size differences between adults and juveniles. The present study concretizes this vision by implementing counting and body area measurement simultaneously within a single AI system, enabling automated derivation of the standard endpoints while also providing a quantitative basis for novel body-area-based endpoints and population-structure-level analysis. In particular, the automated measurement of juvenile body area makes it possible to characterize the size distribution within a population quantitatively, providing a foundation for advancing beyond count-based assessment toward population-level evaluation through changes in cohort structure.

### 4.2. Assessment of population-level toxic responses through cohort analysis

Because the cohorts in this study were analyzed at a single time point, it is difficult to determine definitively whether the two statistically distinguished groups represent age groups or merely size groups. However, a time-series study of *A. kimi* reported two age groups over the 28 days following oviposition (Son et al., 2009). In the present study, two cohorts were reproducibly observed across all treatments, and the mean body area of each cohort remained within a similar range regardless of exposure concentration, suggesting that the cohort structure reflects a property inherent to the developmental stages of *A. kimi* rather than an artifact of oviposition pattern or statistical chance. The two cohorts were therefore interpreted as an age-group distribution.

For all metal(loid)s, the proportion of the small cohort consistently increased and that of the large cohort decreased with increasing concentration, with the large cohort nearly disappearing at the highest concentrations. A similar pattern was reported by Crouau and Moïa (2006), who found that the larger modal group of *Folsomia candida* juveniles decreased preferentially with increasing concentration. This shift may arise from two mechanisms. First, younger individuals may have died before developing into the older stage, consistent with the markedly higher sensitivity of juveniles than adults to metal(loid)s (An et al., 2013). Second, even without causing death, exposure may have slowed molting and growth and thereby delayed the transition to the next developmental stage. Because Collembola grow by molting throughout life—advancing through successive instars only as they molt— and because both molting and somatic growth are energetically expensive, the energy that toxicant stress reallocates to detoxification and maintenance is drawn from these processes, lowering molting frequency and growth rate, slowing instar progression, and prolonging residence in the younger, smaller stages (Crommentuijn et al., 1993, 1995; Kristiansen et al., 2021).

In either case, this suggests that the number of juveniles counted in the 28-day reproduction test may be governed more by the survival and development of juveniles during exposure than by the oviposition capacity of the adults. Indeed, several studies have reported that toxicant exposure affects the growth and survival of the juveniles produced without affecting adult oviposition capacity (Hani et al., 2019; Tian et al., 2020; Zheng et al., 2022), and Min et al. (2025) similarly concluded that the 28-day juvenile count reflects the juveniles’ direct exposure to contaminated soil more than the adults’ reproductive capacity.

The present study thus provides empirical support for the proposal that changes in cohort structure constitute a more meaningful endpoint than simple growth or reproduction measures (Marks et al., 2015; Palamara et al., 2022). Both mechanisms outlined above converge on the same outcome—the younger cohort failing to develop into the larger cohort, or doing so more slowly—and this pattern may serve as an indirect indicator of the extent to which the population’s progression toward the next generation is impaired. In this sense, the shift in cohort composition functions as a toxicity endpoint that captures effects at the level of population structure rather than as an average across individuals, not only offering a new perspective for toxicity assessment but also helping to interpret changes in conventional toxicity endpoints in greater depth.

### 4.3. Suitability of body area as a growth indicator and the potential of cumulative body area as a toxicity endpoint

In quantifying individual size, a comparison of body length, body width, and body area showed that body area was as sensitive as, or more sensitive than, the linear dimensions depending on the metal(loid), with this tendency being relatively pronounced for the cumulative endpoints. This likely arises from the nature of body area as a two-dimensional measure integrating body length and width, which appears to reflect the three-dimensional body form and body weight of Collembola somewhat better than a single linear dimension, in line with reports of a high correlation between body weight and body area in invertebrates (Wardhaugh, 2013; Gomes et al., 2025). In addition, body length is prone to measurement error from the curvature of individuals—prompting correction methods that exclude curved specimens (Bánszegi et al., 2014)—so body area is considered to offer certain practical advantages in an image-based automated environment.

Among the three endpoints, cumulative body area was generally the most sensitive, whereas mean body area tended to be relatively less sensitive, a pattern that may be explained by the influence of cohort structure. As shown by the preceding cohort analysis, the mean body area of juveniles appears to be governed more by the declining proportion of the large cohort than by growth inhibition of individual organisms, and therefore tends to be less sensitive than the number of individuals. This is consistent with previous assessments in which growth indicators have been treated as auxiliary endpoints to mortality and reproduction (Crouau and Moïa, 2006; Bur et al., 2010).

In the case of Cu, an exceptional pattern was observed in which the mean body area decreased significantly somewhat earlier than the number of individuals. Cu is an essential nutrient that induces hormesis at low concentrations (Ardestani and van Gestel, 2013a; Ardestani et al., 2013b); here, the number of individuals first increased and then decreased owing to hormesis, whereas the mean body area decreased gradually without such an increase, thus appearing to reflect the toxic effect relatively early. This difference was even clearer at the cohort level, where the small cohort showed a hormetic increase while the large cohort decreased progressively with increasing Cu concentration. This cohort-level difference can be reconciled with the proposal of Jager et al. (2013) that hormesis-driven increases in oviposition may yield underdeveloped juveniles, and with the finding of Tyne et al. (2015) that low-concentration exposure elicits hormesis in already-developed adults while exerting toxicity during prolonged exposure from the juvenile stage—consistent with our interpretation that the small cohort is influenced more by adult oviposition and the large cohort more by subsequent survival and growth.

Cumulative body area emerged as a relatively consistently sensitive indicator because it simultaneously reflects the number of individuals and individual size. In ecotoxicity assessment, individual endpoints such as survival, growth, and reproduction each provide independent information, but evaluating them in isolation may not sufficiently capture the physiological burden that chemicals impose. In a similar vein, an integrative indicator combining individual size and abundance—body area multiplied by the number of individuals—has recently been reported as the most sensitive endpoint in a *Folsomia candida* toxicity test (Gomes et al., 2025). In a study on *Daphnia*, Bosker et al. (2019) further proposed such cumulative biomass as a means of extending the assessment to an indicator of population-level change.

The present results also appear to be consistent with this view. The juvenile response was driven by shifts in cohort composition, so the mean body area was diluted by the changing proportion of the two cohorts and the count alone tended to reflect only part of the change. By contrast, cumulative body area, integrating abundance and individual size, responded comparatively strongly and emerged as the most sensitive endpoint under the present test conditions. In this respect, cohort analysis and cumulative body area may be used in a complementary manner: the former provides clues for inferring how the juvenile population is restructured, whereas the latter offers a single-value indicator that sensitively captures that change.

AI-based image analysis enabled automated measurement of body length, body width, and body area for all individuals rather than a subset, thereby making cumulative body area—which was difficult to obtain manually—available as a measurable endpoint. Moreover, area data for all individuals allow body-area distribution and cohort structure (large and small cohorts) to be characterized quantitatively, beyond simple mean values, which can help track how the juvenile population is restructured under exposure. On this measurement basis, cumulative body area integrates abundance and individual size into a single value, while cohort analysis explains the structural background of that change, allowing the two to operate complementarily. Consequently, the framework combining these two approaches can contribute to interpreting not only whether a toxic effect occurs but also how that effect influences population structure, and is considered to help complement conventional individual endpoints in capturing sublethal toxicity in a more sensitive and ecologically meaningful way.

More broadly, these results indicate that AI-enabled, population-wide measurement can add endpoints that are at once more sensitive and more ecologically interpretable than abundance alone. Because the integrative endpoints yield lower effect concentrations, incorporating them could lead to more protective soil quality criteria, directly linking the methodological advance to the regulatory goal of soil protection. The framework is, in principle, transferable: just as automated counting has been validated across five laboratories for *Folsomia candida* (Wehrli et al., 2026), the body-area and cohort endpoints introduced here could be extended to other standard test species and to multi-laboratory ring tests, providing a concrete route toward incorporating population-structure endpoints into the OECD 232 and ISO 11267 protocols.

## 5. Conclusion

Since 1984, the OECD soil quality test guidelines have sought to establish biologically meaningful effect thresholds that protect soil ecosystem functions from chemical contamination (OECD, 1984), progressively refined around sensitive, directly exposed bioindicator species such as Collembola (ISO, 1999, 2014, 2023). Within this framework, however, the effect concentrations underpinning soil quality criteria have been derived almost entirely from abundance-based endpoints, because, in the pre-AI era, measuring growth and population structure across all individuals— although its value had been recognized—was so labor-intensive that it was rarely undertaken (Gruss et al., 2022; Suzuki et al., 2015). By making such population-wide measurement straightforward, the present system shows that more sensitive, integrative endpoints—cumulative body area and juvenile cohort structure—yield consistently lower effect concentrations than the standard count. This raises the possibility that thresholds derived from abundance alone may underestimate the toxicity of soil contaminants, leaving current criteria less protective of soil ecosystem function than intended.

Although based on a single species and three metal(loid)s and requiring broader validation, these findings provide a basic technical and scientific basis for re-establishing the relationship between sensitive bioindicators and contaminants, and ultimately for developing more stringent, ecologically protective soil quality criteria.

## CRediT authorship contribution statement

**Hyun-Gi Min:** Data curation, Visualization, Writing – original draft. **Lee-Hyeon Jeon:** Investigation, Formal analysis, Data curation. **Jeongwon Choi:** Investigation, Validation, Formal analysis. **Han Soo Kim:** Investigation, Resources, Writing – review & editing. **June Wee:** Methodology, Investigation, Visualization, Writing – review & editing. **Wonjun Hong:** Software, Validation, Formal analysis. **Daejin Choi:** Methodology, Software, Validation, Supervision, Project administration, Writing – review & editing. **Yun-Sik Lee:** Conceptualization, Methodology, Supervision, Project administration, Funding acquisition, Writing – review & editing.

## Data availability

Data will be made available on request, and the code for the AI-based detection and body-area measurement is openly accessible at https://github.com/HAI-at-Ewha/alkimi-size.

## Declaration of generative AI and AI-assisted technologies in the manuscript preparation process

During the preparation of this work the authors used Claude (Anthropic) in order to improve the language and readability of the manuscript. After using this tool, the authors reviewed and edited the content as needed and take full responsibility for the content of the published article.

