## Supplementary Material (Fig. S1, Tables S1-S7) for "Deep learning-based counting and body-area measurement enable sensitive population-level endpoints in the Collembola reproduction test"

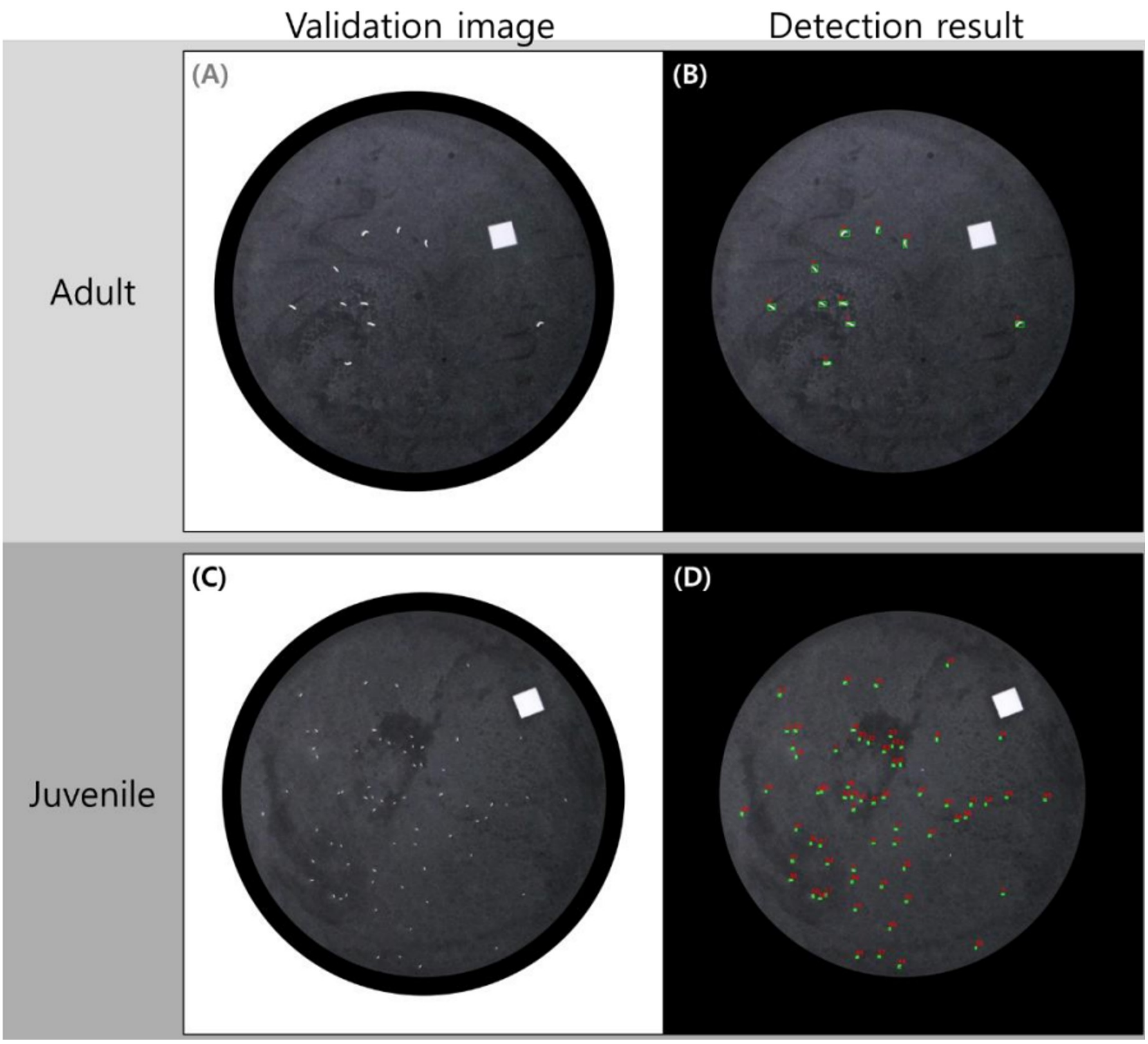


**Fig. S1.** Representative validation images of *Allonychiurus kimi* (A, C) and the corresponding detection results of the AI-based model (B, D) for adults (A, B) and juveniles (C, D). In the detection results, each detected individual is enclosed in a bounding box. The white square (5 mm × 5 mm) on each substrate is the marker used for pixel-to-area conversion.

**Table S1.** Concentration-dependent agreement between manual and AI-based body area measurements of *Allonychiurus kimi* exposed to cadmium (Cd).

| Developmental stage | Cd concentration (mg $kg^{-1}$) | *r*RMSE[a] | $R^{2b}$ | Body area ($mm^2$) | |
|---|---|---|---|---|---|
| | | | | Manual | Automatic |
| Adult | 0 | 1.14 | 1.00 | 0.672 ± 0.180 | 0.671 ± 0.177 |
| | 12.5 | 1.55 | 0.99 | 0.543 ± 0.118 | 0.541 ± 0.117 |
| | 25 | 2.34 | 0.99 | 0.442 ± 0.112 | 0.443 ± 0.111 |
| | 50 | 6.14 | 0.97 | 0.283 ± 0.099 | 0.290 ± 0.100 |
| | 100 | 3.93 | 0.98 | 0.277 ± 0.069 | 0.286 ± 0.067 |
| | 200 | 4.57 | 0.93 | 0.314 ± 0.053 | 0.323 ± 0.049 |
| Juvenile | 0 | 4.61 | 0.99 | 0.107 ± 0.044 | 0.109 ± 0.045 |
| | 12.5 | 6.64 | 0.98 | 0.079 ± 0.038 | 0.081 ± 0.036 |
| | 25 | 15.09 | 0.84 | 0.076 ± 0.029 | 0.078 ± 0.029 |
| | 50 | 6.37 | 0.98 | 0.069 ± 0.033 | 0.071 ± 0.033 |
| | 100 | 7.12 | 1.00 | 0.054 ± 0.060 | 0.053 ± 0.059 |
| | 200 | 4.80 | 0.86 | 0.064 ± 0.008 | 0.066 ± 0.010 |

Values are the mean ± SD across all individuals measured at each concentration; SD, standard deviation.

[a] Relative root mean square error.

[b] Coefficient of determination relative to the 1:1 line

**Table S2.** Concentration-dependent agreement between manual and AI-based body area measurements of *Allonychiurus kimi* exposed to arsenic (As).

| Developmental stage | As concentration (mg kg$^{-1}$) | *r*RMSE[a] | $R^{2}$[b] | Body area (mm$^2$) | |
|---|---|---|---|---|---|
| | | | | Manual | Automatic |
| Adult | 0 | 1.44 | 0.99 | 0.584 ± 0.101 | 0.589 ± 0.101 |
| | 10 | 1.98 | 0.99 | 0.570 ± 0.139 | 0.578 ± 0.142 |
| | 20 | 1.80 | 1.00 | 0.490 ± 0.140 | 0.496 ± 0.139 |
| | 40 | 1.23 | 1.00 | 0.353 ± 0.093 | 0.358 ± 0.092 |
| Juvenile | 0 | 8.92 | 0.91 | 0.092 ± 0.028 | 0.094 ± 0.026 |
| | 10 | 11.66 | 0.82 | 0.073 ± 0.020 | 0.078 ± 0.018 |
| | 20 | 8.09 | 0.99 | 0.075 ± 0.019 | 0.079 ± 0.017 |
| | 40 | – | – | – | – |

Values are the mean ± SD across all individuals measured at each concentration; SD, standard deviation. No juveniles were recovered at 40 mg kg$^{-1}$.

[a] Relative root mean square error.

[b] Coefficient of determination relative to the 1:1 line.

**Table S3.** Concentration-dependent agreement between manual and AI-based body area measurements of *Allonychiurus kimi* exposed to copper (Cu).

| Developmental stage | Cu concentration (mg $kg^{-1}$) | *r*RMSE[a] | $R^{2b}$ | Body area ($mm^2$) | |
|---|---|---|---|---|---|
| | | | | Manual | Automatic |
| Adult | 0 | 1.38 | 1.00 | 0.744 ± 0.167 | 0.740 ± 0.166 |
| | 50 | 1.31 | 0.99 | 0.711 ± 0.130 | 0.705 ± 0.127 |
| | 100 | 2.15 | 0.99 | 0.810 ± 0.143 | 0.797 ± 0.136 |
| | 200 | 1.65 | 0.99 | 0.724 ± 0.113 | 0.716 ± 0.109 |
| | 400 | 1.14 | 0.99 | 0.629 ± 0.087 | 0.626 ± 0.088 |
| | 800 | 1.17 | 1.00 | 0.534 ± 0.122 | 0.532 ± 0.120 |
| Juvenile | 0 | 3.13 | 0.99 | 0.209 ± 0.062 | 0.212 ± 0.063 |
| | 50 | 3.99 | 0.98 | 0.175 ± 0.052 | 0.177 ± 0.052 |
| | 100 | 8.86 | 0.95 | 0.209 ± 0.080 | 0.209 ± 0.078 |
| | 200 | 7.83 | 0.97 | 0.182 ± 0.083 | 0.183 ± 0.080 |
| | 400 | 3.83 | 0.99 | 0.132 ± 0.043 | 0.136 ± 0.044 |
| | 800 | 3.40 | 0.98 | 0.127 ± 0.034 | 0.130 ± 0.034 |

Values are the mean ± SD across all individuals measured at each concentration; SD, standard deviation.
[a] Relative root mean square error.
[b] Coefficient of determination relative to the 1:1 line.

**Table S4.** Determination of the number of cohorts per concentration based on ΔBIC and bootstrap LRT

| Substance | Concentation (mg kg$^{-1}$) | N[a] | Number of cohorts | ΔBIC[b] | Bootstrap LRT[c] (1 vs 2), p |
|---|---|---|---|---|---|
| Cd | 0 | 348 | 2 | 66.82 | <0.0001 |
| | 12.5 | 256 | 2 | 23.99 | <0.0001 |
| | 25 | 140 | 2 | 15.08 | <0.0001 |
| | 50 | 69 | 2 | 18.90 | 0.0002 |
| As | 0 | 254 | 2 | 57.24 | <0.0001 |
| | 10 | 149 | 2 | 11.42 | <0.0001 |
| | 20 | 107 | 1 | — | — |
| Cu | 0 | 292 | 2 | 3.75 | 0.0006 |
| | 50 | 315 | 2 | 2.19 | 0.0004 |
| | 100 | 353 | 2 | 32.73 | <0.0001 |
| | 200 | 383 | 2 | 151.39 | <0.0001 |
| | 400 | 227 | 2 | 8.05 | <0.0001 |
| | 800 | 64 | 1 | — | — |

For single-cohort treatments, ΔBIC and the LRT *p*-value are reported as "—".
[a] Number of juveniles analyzed.
[b] Difference in Bayesian Information Criterion between the two- and one-cohort models; following Kass and Raftery (1995), ΔBIC ≥ 2, > 6, and > 10 indicate positive, strong, and very strong evidence, respectively.
[c] Bootstrap likelihood-ratio test of one versus two cohorts (mclustBootstrapLRT, "V" model, 9999 replications).

**Table S5.** $EC_{50}$ values for juvenile *Allonychiurus kimi* endpoints based on manual measurements following exposure to cadmium (Cd), arsenic (As), and copper (Cu).

| Endpoint | Cd | As | Cu |
|---|---|---|---|
| Average body area | 58.0 (42.0–80.1) | 24.0 (2.2–194.1) | 1137.9 (591.5–2188.4) |
| Average body length | 58.6 (6.3–484.8) | 23.8 (N.C.) | 5772.5 (1345.5–24754.7) |
| Average body width | 58.7 (29.9–114.2) | 24.3 (N.C.) | 2595.6 (778.8–8645.8) |
| Cumulative body area | 13.7 (9.4–20.0) | 9.8 (5.6–16.7) | 394.0 (317.2–489.4) |
| Cumulative body length | 17.7 (14.7–21.3) | 11.7 (7.4–18.1) | 453.3 (346.5–593.0) |
| Cumulative body width | 15.8 (12.1–20.4) | 11.4 (7.2–17.7) | 431.6 (336.6–553.3) |

Values are $EC_{50}$ with 95 % confidence intervals in parentheses, expressed in mg $kg^{-1}$. N.C., not calculable. Average and cumulative endpoints denote the per-individual mean and the summed total per treatment, respectively, for body area, body length, and body width.

**Table S6.** No-observed-effect concentration (NOEC) and lowest-observed-effect concentration (LOEC) for three endpoints (number of individuals, average body area, and cumulative body area) of *Allonychiurus kimi* adults and juveniles exposed to three metal(loid)s (Cd, As, Cu) for 28 days.

| Test substance | Developmental stage | Endpoint | NOEC (mg $kg^{-1}$) | LOEC (mg $kg^{-1}$) |
|---|---|---|---|---|
| Cd | Adult | Number of individuals | 12.5 | 25 |
| | | Average body area | 12.5 | 25 |
| | | Cumulative body area | < 12.5 | 12.5 |
| | Juvenile | Number of individuals | < 12.5 | 12.5 |
| | | Average body area | 50 | 100 |
| | | Cumulative body area | < 12.5 | 12.5 |
| As | Adult | Number of individuals | 10 | 20 |
| | | Average body area | 20 | 40 |
| | | Cumulative body area | 10 | 20 |
| | Juvenile | Number of individuals | < 10 | 10 |
| | | Average body area | 20 | 40 |
| | | Cumulative body area | < 10 | 10 |
| Cu | Adult | Number of individuals | 400 | 800 |
| | | Average body area | 400 | 800 |
| | | Cumulative body area | 400 | 800 |
| | Juvenile | Number of individuals | 400 | 800 |
| | | Average body area | 200 | 400 |
| | | Cumulative body area | 200 | 400 |

NOEC and LOEC were determined by one-way ANOVA followed by Dunnett’s test ($p < 0.05$). A NOEC below the lowest tested concentration is reported as “< ” followed by that concentration.

**Table S7.** Pairwise comparison of $EC_{50}$ values among the three endpoints by Z-test for *Allonychiurus kimi* exposed to cadmium (Cd), arsenic (As), and copper (Cu).

| Test substance | Comparison | $EC_{50}$[a] (1) | $EC_{50}$ (2) | z | *p*-value |
|---|---|---|---|---|---|
| Cd | Number of individuals vs Average body area | 21.09 | 65.28 | −2.731 | 0.006 |
| | Number of individuals vs Cumulative body area | 21.09 | 14.10 | 1.954 | 0.051 |
| | Average body area vs Cumulative body area | 65.28 | 14.10 | 3.484 | 0.001 |
| As | Number of individuals vs Average body area | 13.60 | 23.87 | −0.544 | 0.586 |
| | Number of individuals vs Cumulative body area | 13.60 | 10.53 | 0.780 | 0.435 |
| | Average body area vs Cumulative body area | 23.87 | 10.53 | 0.780 | 0.436 |
| Cu | Number of individuals vs Average body area | 513.72 | 1223.36 | −2.233 | 0.026 |
| | Number of individuals vs Cumulative body area | 513.72 | 398.80 | 1.346 | 0.178 |
| | Average body area vs Cumulative body area | 1223.36 | 398.80 | 3.011 | 0.003 |

$EC_{50}$ (1) and $EC_{50}$ (2) are the $EC_{50}$ values (mg $kg^{-1}$) of the first and second endpoints in each comparison, respectively. The Z statistic and *p*-value test the difference between the two $EC_{50}$ estimates; $p < 0.05$ indicates a significant difference. "Number of individuals" denotes the number of juveniles.

[a] Effective concentration required to reduce the endpoint by 50 %.
